# runibic: a Bioconductor package for parallel row-based biclustering of gene expression data

**DOI:** 10.1101/210682

**Authors:** Patryk Orzechowski, Artur Pańszczyk, Xiuzhen Huang, Jason H. Moore

## Abstract

**Motivation:** Biclustering (called also co-clustering) is an unsupervised technique of simultaneous analysis of rows and columns of input matrix. From the first application to gene expression data, multiple algorithms have been proposed. Only a handful of them were able to provide accurate results and were fast enough to be suitable for large-scale genomic datasets.

**Results:** In this paper we introduce a Bioconductor package with parallel version of UniBic biclustering algorithm: one of the most accurate biclustering methods that have been developed so far. For the convenience of usage, we have wrapped the algorithm in an R package called *runibic*. The package includes: (1) a couple of times faster parallel version of the original sequential algorithm,(2) muchmore efficient memory management, (3) modularity which allows to build new methods on top of the provided one, (4) integration with the modern Bioconductor packages such as *SummarizedExperiment*, *ExpressionSet*and *biclust*.

**Availability:** The package is implemented in R (3.4) and will be available in the new release of Bioconductor (3.6). Currently it could be downloaded from the following URL: http://github.com/athril/runibic/

**Supplementary information:** Supplementary informations are available in vignette of the package.

## 1. Introduction

The recent advantages in transcriptomic analysis, including development of high-throughput and high-resolution platforms including RNA-seq, single-cell RNA-sequencing (scRNA-seq) or high-throughput PCR have allowed to design experiments that provide datasets with even hundreds of thousands columns and thousands rows. This have set new requirements for data analytics. Modern methods need to yield accurate results. They are required to handle large datasets and are expected to finish computations in reasonable time.

With growing amount of genomic data there is an urgent need for efficient and precise methods that are able to capture the underlying patterns in gene expression datasets. One of the techniques that proved to be very insightful in gene expression analysis is biclustering, which allows to detect subsets of genes and samples in complex and noisy data. Biclustering is considered NP-hard as it investigates relations between multiple rows that occur in different subsets of columns. The running time of the algorithms is usually highly dependent on the size of the input data.

The vast majority of biclustering methods are sequential. There area couple of common reasons for this. Some methods are specifically designed to yield only one bicluster at a time. Each runofthealgorithm depends on the previous findings. Other methods use graph-based structures, which are difficult to parallelize, or perform hardly scalable statistical analyses. For some group of the methods parallel implementation may not be beneficial, as they extensively use binary operations. Bioconductor in version 3.5 provides the following biclustering methods and packages for gene expression analysis:

- ISA (Bergmann *et al.*, 2003) - implemented in eisa and isa2 Bioconductor packages (Csardi *et al.*, 2010),
- CC (Cheng and Church, 2000), Plaid methods Lazzeroni and Owen (2002), Bimax (Prelic´ *et al.*, 2006), xMotifs (Murali and Kasif, 2003), Quest (Kaiser, 2011), Spectral Kluger *et al.* (2003) - all available in biclust package (Kaiser *et al.*, 2015),
- FABIA, FABIAS, FABIAP - available in Bioconductor package fabia (Hochreiter *et al.*, 2010),
- HapFABIA - implemented in package hapfabia (Hochreiter, 2013)
- QUBIC (Li *et al.*, 2009) - implemented in more modern package QUBIC (Zhang *et al.*, 2017) and package older rqubic (Zhang, 2015),
- MCbiclust - available in Bioconductor package MCbiclust (Bentham, 2017),
- SSVD (Lee *et al.*, 2010) and S4VD (Sill *et al.*, 2011) - available in Bioconductor package s4vd Sill and Kaiser (2015),
- Iterative Binary Biclustering of Gene sets - available in Bioconductor package iBBiG Gusenleitner *et al.*(2012)

The vast majority of the method are implemented in R, which is slower than C. Some of the methods, e.g. QUBIC, benefit from calls to high-performance C++ linear algebra libraries, such as *Rcpp* (Eddelbuettel and François, 2011) and *RcppArmadilo* (Eddelbuettel and Sanderson, 2014). The comparison of R packages functionality is presented in Table 1.

**Table 1.**
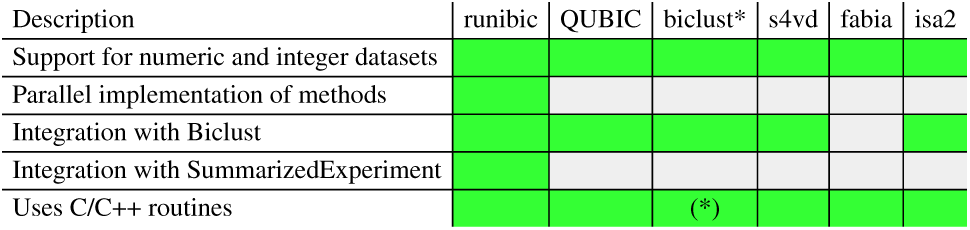
Comparison of functionalities of different R packages. (*) - Only Bimax algorithm uses wrapped C function call.

One of the recent breakthroughs in gene expression analysis was UniBic (Wang *et al.*, 2016). The algorithm originally implemented in C managed to capture biologically meaningful trend-preserving patterns and proved to outperform multiple other methods. The method also showed great potential for parallelization. Unfortunately the implementation of the method wasn’t efficient enough and the code had some memory leaks.

## 2 Methods

In this paper we introduce aBioconductorpackagecalled *runibic* with parallel implementation of one of the most accurate biclustering methods: UniBic. Thealgorithm, orignally released as sequential, has proven to outperform multiple popular biclustering state-of-the-art biclustering methods (Wang *et al.*, 2016). We have redesigned the code and reimplemented the method into more modern C++11 programming language. By parallelizing chunks of the code using OpenMP standard Dagum and Menon (1998), we obtained approximately up to a couple of times speedup in terms of execution time for popular genomic datasets. By fixing some of the memory management bugs, our package provides more stable and reliable implementation of UniBic algorithm.

### 2.1 Implementation

In the provided implementation of UniBic algorithm, we migrated the original code from C to C++11 programming language and added OpenMP support. Code refactoring allowed us to take advantage of multiple aspects of modern-style language programming:

- safer and more modern memory management replaced difficult to maintain C style memory allocations and deallocations,
- fast and efficient containers from Standard Template Library (STL), such as vectors, sets and algorithms, were used for acceleration of common operations like iterate, sort, search, count, or copy,
- the original implementation in most cases allocated a large number of simple arrays and used loops with slow indexing for common operations,
- removing many redundant copying and memory allocations,
- fixing a couple of memory leaks, which caused segmentation fault for some datasets.

Porting thecodeimprovedinterpretabilityofthecodeallowedto remove multiple redundancies present in the previous UniBic implementation. For example, we replaced the original four functions that calculated the Longest Common Subsequence with a single one with multiple options. Similar improvements were made in other code sections, for example: in discretization, in calculation of Longest Common Subsequence between each pair of rows, in clustering and bicluster expansion parts. In order to provide more insightful analysis into the modules of UniBic algorithm, we separated and exported the major steps of the original method. Thus, the algorithm may be run using either a single command, or executed step by step. This provides much better control, improves clarity of the method,and allows its future customization. The algorithm provided in *runibic* package is divided into the following sections:

- *set_runibic_params* - a function that sets parameters for algorithm,
- *runiDiscretize* - the original UniBic discretize approach, which take into account the number of ranks from ‘div’ parameter and quantile value from ‘q’ parameter (Step 1 of the method),
- *unisort* - a function that sorts the rows of a matrix and returns the indexes of sorted columns in each row (Step 2),
- *calculateLCS* - a function that calculates the Longest Common Subsequence (LCS) between each unique pair of rows in the matrix, returns a list of LCS lengths and row pairs (Step 3),
- *cluster* - the main biclustering method which builds biclusters based on the input data and *calculateLCS* results (Steps 4 and 5).

By designing a modular structure of the package we intended to simplify flexible modifications of the original algorithm. Such methods may use different preprocessing or ranking techniques, or expand bicusters using different rows as seeds. An example includes different method of sorting results from *calculateLCS*. The proposed method, which is based on a stable STL sort, could be used as an alternative to the old C style pointer and sorting based on Fibonacci Heap. In our opinion the proposed method is more robust and better reflects the original intention. The choice of the method may implicate the outcome of the algorithm, as different LCSes of the same length may be chosen as seeds.

### 2.2 Parallelization

In order to improve the algorithmexecutiontimethemostcrucialand computationally intensive parts of the code were parallelized using OpenMP standard. One of the most time consuming steps of UniBic is calculating Longest Common Subsequence (LCS) between unique pairs of rows. We rearranged the code and achieved parallelization where each core of the CPU calculates unique LCS between unique pair of rows simultaneously. Similarly, we also paralleled the data preprocessing required by the method, so as expansions of each of the biclusters, which required calculations of LCS between each row and the seed. All mentioned operations allowed us to obtain biclustering results in several minutes on the modern computer with modern processor.

### 2.3 Integration with Bioconductor packages

The *runibic* package takes advantage of *Rcpp* library that allows seamless integration of C++ code with R environment. The *runibic* package is also integrated with *biclust* package methods for biclustering process. Results returned from *runibic* are wrapped into a *Biclust* object, which can be used for further examination, including visualization and analysis provided by *biclust* package. The examples of usage are presented in Supplementary material as well as in the package manual.

~~~
library (runibic)
library (biclust)
test <− matrix (rnorm(1000), 100, 100)
res <− runibic (test)
res <− biclust :: biclust (test, method = BCUnibic())
~~~

Similarly, the biclust method could be applied to any matrix extracted from *ExpressionSet* using *exprs*() function.

### 2.4 Support for SummarizedExperiment

Apart from allowing analysis of genomic data from historical *ExpressionSet*, *runibic* package is compatible with *SummarizedExperiment* class (Morgan *et al.*, 2017). This class offers much more flexibility in terms of experiment design and supports both RNA-Seq and Chip-Seq. This makes *runibic* a very easy tool for performing biclustering analysis of modern genomic experiments. An example of using *runibic* with Single-Cell RNA-Seq Datasets is provided in Supplementary material.

## 3 Results

To investigate running times of the method, we have applied it to several popular datasets. The running times of the revised and the original UniBic algorithm as well as the revised parallel version are presented in Table 2.

**Table 2.**
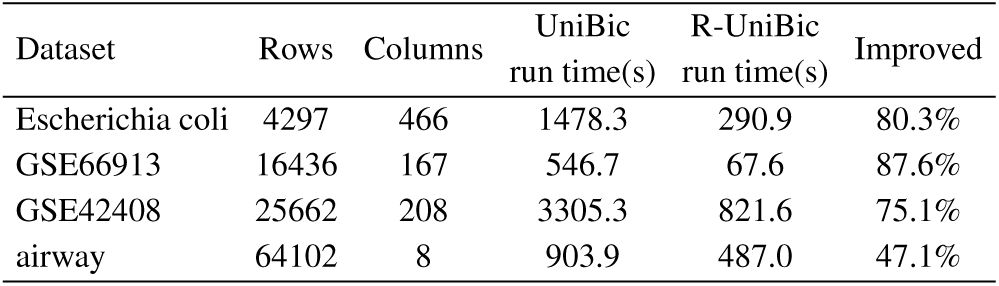
Running times of the original version of UniBic Wang et al. (2016) and parallel UniBic in R from Bioconductor package.

By refactoring and optimizing the code,the new parallel version of UniBic biclustering algorithm provided by *runibic* package is approximately 2-5 times faster than the original version of the algorithm for popular genomic datasets.

## 4 Conclusions

In this paper we introduce *runibic* package with revised and parallelized version of UniBic algorithm. The package is going to be available in the newest Bioconductor 3.6 release.. Providing a modular structure of the package allows to easily understand steps of the method and makes code much more interpretable. The *runibic* package provide *runibic* method that could be applied to any matrix in R, expression set extracted from *ExpressionSet* or *SummarizedExperiment* class. Integration with many common R and Bioconductor packages (e.g. *biclust*, *QUBIC*), as well as extensive documentation on one of the most accurate biclustering methods developed so far, make *runibic* package easily accessible and very flexible for gene expression analysis.

## 5 Funding

This research was supported in part by PL-Grid Infrastructure and by grant LM012601 from the National Institutes of Health (USA).

